# Comparative Connectomics Highlights Conserved Architectural Synaptic Motifs in the *Drosophila* Mushroom Body

**DOI:** 10.1101/2025.09.22.677863

**Authors:** Patricia K. Rivlin, Michael Robinette, Jordan K. Matelsky, Brock Wester

**Affiliations:** Johns Hopkins Applied Physics Laboratory. Laurel, MD, USA; Department of Bioengineering, University of Pennsylvania. Philadelphia, PA, USA

## Abstract

While the influence of synaptic plasticity on learning and memory has been extensively studied, the detailed patterns of synaptic connectivity remain incompletely mapped. Convergent synaptic motifs — a tight grouping of at least two axons whose active zones are within 300nm and which are presynaptic to the same target — are a common feature of neural circuits in the insect brain and are believed to serve as an important computational primitive in many brain areas. The Mushroom Body (MB) of *Drosophila*, for instance, is the center of associative learning and memory, where sensory information is conducted by Kenyon cells (KCs), the intrinsic neurons of the MB, and integrated by MB output neurons (MBONs). Indeed, the majority of KC-to-MBON synapses occur in a convergent motif. Nonetheless, the functional role of this convergent motif is not well studied. To gain insight into their potential role in the MB, we combine big-data network neuroscience tools with existing electron microscopy connectome datasets to detect and map the distribution of convergent synaptic motifs. We find that convergent motifs consistently occur across the MB in different individuals, including the *α*-lobe where they were first quantified, and we report on both the variance and consistency in the formation of these motifs across different MB regions and individuals. Our discovery of multiply-convergent motifs — where two KCs target multiple postsynaptic targets simultaneously — reveals a previously unrecognized synaptic economy that may optimize information transfer while conserving neural resources. These stereotyped arrangements likely represent fundamental organizational principles underlying associative learning across species. Lastly, to our knowledge, this study offers the first and most extensive comparative analysis of synaptic motifs across *Drosophila* connectomes, establishing a framework for enabling systematic motif analysis of synapses across species.

## 1 Introduction

Synapses exhibit structural diversity in both pre- and post-synaptic arrangements across different species and brain regions. Often, synapses form local motifs — repeating substructures that the brain reuses with only minor modifications. These synaptic motifs remain largely unexplored due to the difficulty of finding and characterizing them. Nonetheless, it is believed that such motifs are crucial for core functions of the brain.^2,3^ Investigating the distribution and cell specificity of these diverse synaptic motifs could offer valuable insights into their functional roles.

Synapses between specific pairs of neuron types can also be stereotyped, consistently exhibiting a particular structural pattern or motif.^4^ For instance, in the *Drosophila* olfactory system, OSN, *µ*PN, and MGN axons form polyadic synapses that differ in the average number of post-synaptic partners, 6, 5, and 3, respectively.^5^ Similarly, in the mouse retina, cone bipolar cell axons form dyads, triads, and tetrads, which vary in their presynaptic configurations (e.g. single versus multiple ribbon), resulting in six distinct synaptic motifs.^4^

Synaptic motifs have been identified at the electron microscopic level in limited subvolumes of the vertebrate and invertebrate brain.^2,4,6–10^ While large-scale connectomes have emerged where millions of pre- and post-synaptic sites are annotated and available for analysis, such connectomes have yet to be leveraged to compare synaptic motifs within and across brain regions and individuals. This may be due, in part, to the level of neuron reconstruction and accuracy of synapse detection in some connectome volumes, as well as the compute cost to analyze synaptic motifs at scale.

The potential importance of convergent motifs is underscored by their appearance across species (e.g., *Drosophila*, locust, octopus) in the learning center as well as the central complex or navigation center of the brain,^11,12^ and analogously as multi-innervated spines in the mammalian brain.^13–15^ As a first demonstration of synaptic motif analysis across individuals and brain subregions, we compare the distribution of convergent synaptic motifs, also called rosettes, across individuals and subregions of the *Drosophila* Mushroom Body (MB).

Three comprehensive connectomes have been generated with EM to study neural circuitry in the *Drosophila* MB: MB-6, centered on the *α*-lobe of the MB;^16^ Hemibrain, a partial brain volume containing the entire MB of the right hemisphere;^1,17^and FlyWire, a whole brain volume.^18^

The Drosophila MB is comprised of ∼2000 intrinsic neurons called Kenyon cells (KCs) whose parallel axons divide the MB into five subregions or lobes, *α, β, α*’, *β* ‘ and *γ*, and form synaptic connections with 20 typical and 14 atypical types of MB output neurons (MBONs) whose dendrites further divide the lobes into compartments.^19^

At the EM level, Drosophila synapses are characterized by a presynaptic process bearing an electron-dense T-bar ribbon, with associated postsynaptic neurites positioned adjacent to the T-bar.^20^ In the fly, central synapses have been most thoroughly investigated in the visual system, where most synapses exhibit a polyadic motif comprising two or more postsynaptic elements associated with a single T-bar. In the MB-6 dataset, KC axons were observed to form three types of synaptic motifs: monadic, polyadic (**Fig. 8** methods), and convergent. Convergent motifs occur when two KC axons (with closely apposed presynaptic sites) converge onto the same dendritic site thus forming a many-to-one configuration (**Fig. 1**). Quantification in the *α* lobe of MB-6 further revealed that the majority of KC-to-MBON synapses (80-93% depending on the MBON type) occur in a convergent motif.

**Figure 1.**
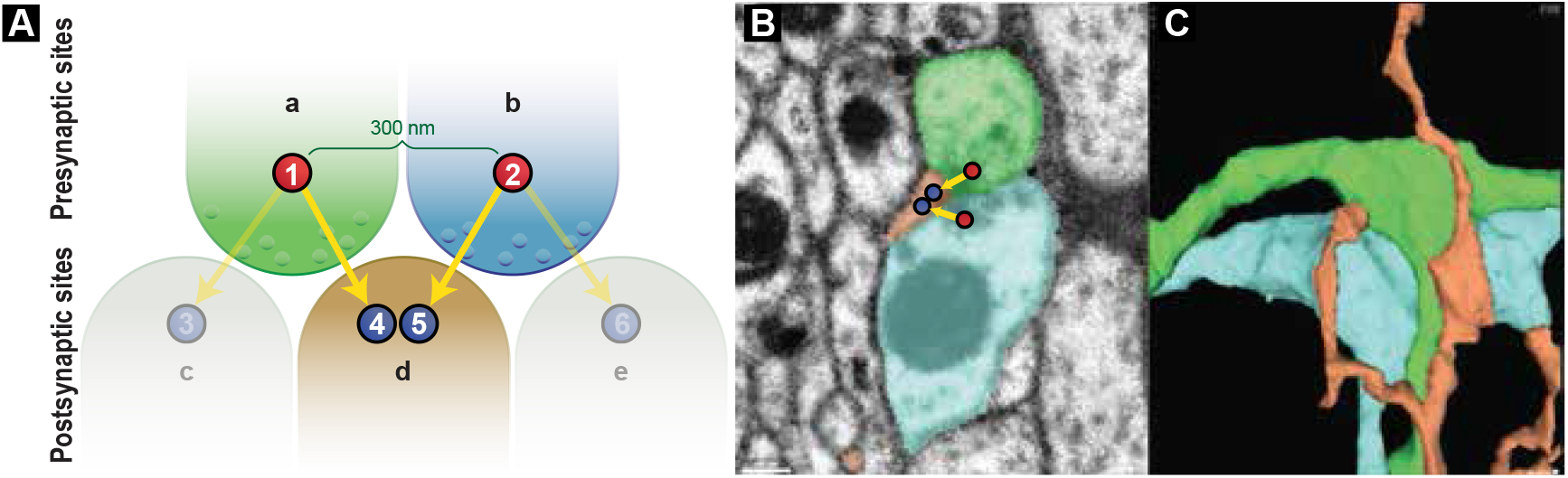
The convergent motif schema and example. **A**. The convergent motif is a common but understudied structure where two very nearby presynaptic sites synapse on the same downstream target postsynaptic site. In this schematic, **a** and **b** represent two distinct presynaptic point annotations that are separated by no more than three hundred nanometers. **B**. A slice of electron microscopy^1^ showing the neuropil that underly this motif (color-coded). **C.** 3D reconstruction reveals the complex structures involved in this motif instance.

Here, we present the first large-scale detection and comparison of synaptic motifs across different brain regions and individuals. We asked whether the rate of convergence is conserved across individuals and MB lobes by analyzing the rate of synaptic convergence in the MB of the Hemibrain and comparing that to the published rate for the *α* lobe of MB-6. During this analysis, we discovered the occurrence of “multiply” convergent motifs, which prompted us to quantify the proportion of single, double and triple motifs formed by each MBON. Given the diversity of convergent motifs, we also asked whether KCs form stereotypic motifs with specific partner types (e.g. MBON types), as previously observed for cone bipolar neurons in the mammalian retina.^4^ Finally, given that higher polyadicity has been correlated with larger synapse size, we investigated whether double convergent synapses are associated with greater polyadicity. Such a relationship would suggest that multiply convergent motifs provide a resource-efficient mechanism for strengthening KC-to-MBON connections while conserving space.

## 2 Results

### 2.1 KC-MBON synapse counts are consistent across animals

An important quality control for connectivity comparisons is the consistency of pre- and post-synapse counts within cell types, both within brains and across brains.^21^ As reported in MB-6, KCs form convergent motifs with every postsynaptic cell type in the *α* lobe, but the fraction of KC synapses to a given cell type that occur in convergent motifs varied between cell types, with highest percentage observed for MBONs (80-93% depending on the MBON type).^16^

In this study, we asked if the percentage of convergent motifs formed by KC-MBON synapses (% convergence) is conserved across individuals and MB lobes. To demonstrate the suitability of MB-6 and hemibrain for these comparisons, we first evaluated the consistency of KC-MBON synapse counts by MBON type in the *α* lobe of MB-6 and hemibrain (**Fig. 2**).

**Figure 2.**
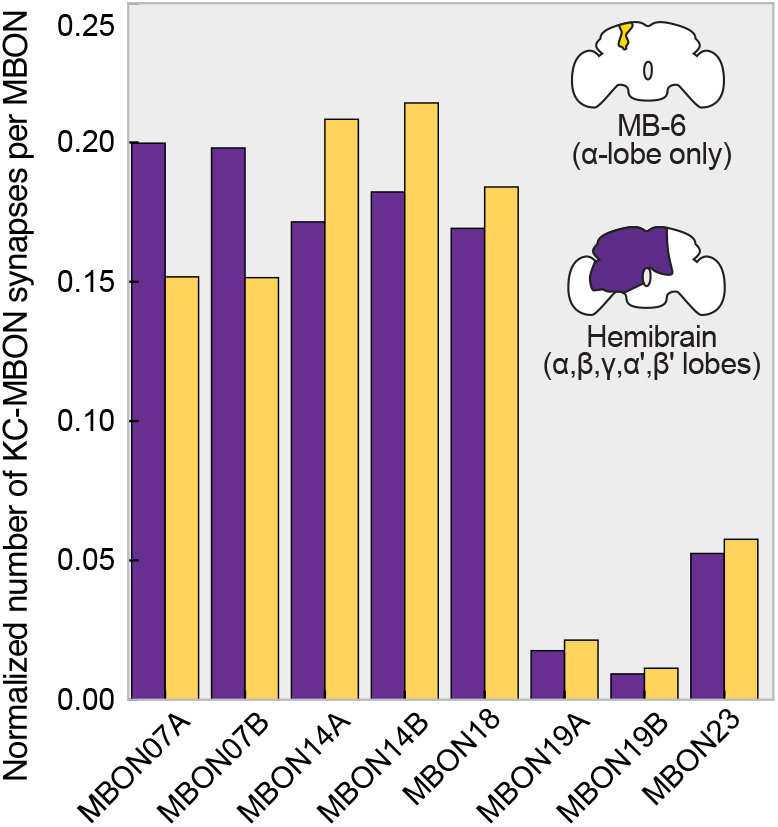
Consistency of KC-MBON synapses across datasets. KC-MBON synapse counts per MBON type in the *α* lobe were normalized to total KC-MBON synapses per dataset.

In MB-6, Takemura et al.^16^ reported that, within each *α*-lobe compartment, every KC axon forms multiple synapses with each MBON, although the average number of KC-MBON synapses per KC varies per MBON type (**Table 1**). To account for differences in the number of KCs innervating each MBON, we also compared the mean number of KC-MBON synapses per KC for each MBON type within and across datasets.

**Table 1.** Comparison of KC→ MBON synaptic connectivity across the MB-6 and hemibrain datasets. See **Fig. 5** for visualization. *: A close examination of MBON07A/B in MB-6 reveals an error in neuPrint, which under-reports the number of MBON07 synapses compared to Takemura et al. (2017). See *Datasets & selection* for more information.

| Dataset | Postsynaptic MBON | Number of pre-synaptic KCs | Total number of KC→MBON synapses | Mean number of KC→MBON synapses per KC |
| --- | --- | --- | --- | --- |
| MB-6 hemibrain | MBON-07A* | 949<br>889 | 9303<br>12830 | 9.8<br>14.43 |
| MB-6 hemibrain | MBON-07B* | 949<br>889 | 9286<br>12720 | 9.79<br>14.31 |
| MB-6 hemibrain | MBON-14A | 948<br>888 | 12770<br>11015 | 13.47<br>12.40 |
| MB-6 hemibrain | MBON-14B | 948<br>887 | 13129<br>11705 | 13.85<br>13.20 |
| MB-6 hemibrain | MBON-18 | 909<br>868 | 11281<br>10867 | 12.41<br>12.53 |
| MB-6 hemibrain | MBON-19A | 236<br>196 | 1311<br>1131 | 5.56<br>5.77 |
| MB-6 hemibrain | MBON-19B | 168<br>223 | 692<br>594 | 4.12<br>2.66 |
| MB-6 hemibrain | MBON-X/23 | 823<br>747 | 3529<br>3371 | 4.29<br>4.51 |

Given that MBON07, MBON14 and MBON19 groups each comprise two neurons (denoted as A and B), we first asked if synapse counts are conserved across neurons of the same MBON type. Within MB-6 and hemibrain, synapse counts and their distribution per KC are consistent across neurons within MBON07 and MBON14, but not MBON19, thus suggesting that MBON19A and MBON19B are reconstructed to different levels of completeness or perhaps even represent different subtypes. Distinguishing between these possibilities would require morphological, connectivity, and molecular analyses (see Discussion).

While we observed consistency in the number of KC-MBON07 and KC-MBON14 synapses within brains, we observed variation across brains, with 37% more KC-MBON07 synapses and 13% fewer KC-MBON14 synapses in the hemibrain as compared to MB-6.

We note that the *α* lobe of MB-6 and the hemibrain were reconstructed to 86% and 84% completeness, respectively,,^1,16^ suggesting that the difference in KC-MBON07 synapse counts is biological (e.g., sexual dimorphism), and the difference in KC-MBON14 counts is due to technical noise.

Schlegel et al.^21^ reported that variations in edge weights of approximately 30% or less could be attributed entirely to technical noise and should be interpreted cautiously when comparing connectivity within and across FlyWire and hemibrain datasets. We anticipate that true biological variance between the male MB-6 dataset and the female hemibrain dataset is low, as both used identical fly rearing conditions, a similar EM staining protocol,^22^ and were imaged with FIB-SEM at the same imaging facility. However, there are differences in synapse detection methods: the hemibrain dataset employed automated detection for both pre- and post-synapses, whereas the MB-6 dataset used automated pre-synapse detection and manual post-synapse detection. Accordingly, with the exception of MBON07, we observed no more than 14% difference in KC-MBON edge weights when comparing MBON types between MB-6 and hemibrain, thus suggesting that these datasets are well suited for comparative motif analysis.

### 2.2 KC-MBON convergence by cell type is conserved across individuals

Takemura et al.^16^ reported that 80–93% of KC-MBON connections (depending on the MBON type) occur in convergent motifs. We asked if the % convergent motifs formed per MBON type is conserved across individuals. Using DotMotif,^2^ we analyzed the proportion of KC-MBON synapses in convergent motifs (hereafter referred to as “% convergence”) in the *α*-lobe of the hemibrain and MB-6. While our analysis confirmed that most KC-MBON synapses occur in convergent motifs in MB-6, we observed a slightly lower % convergence (72–85%) than reported by Takemura et al. This discrepancy may be due to their method of computing convergence by MBON type, which can overestimate % convergence and introduce false positives, as previously discussed. Interestingly, we also observed a slightly lower % convergence in the hemibrain (63–70% per MBON type) as compared to MB-6. However, the trend across MBON types was consistent across datasets, with MBON-14 showing the highest % convergence and MBON-19 the lowest (**Fig. 3**).

**Figure 3.**
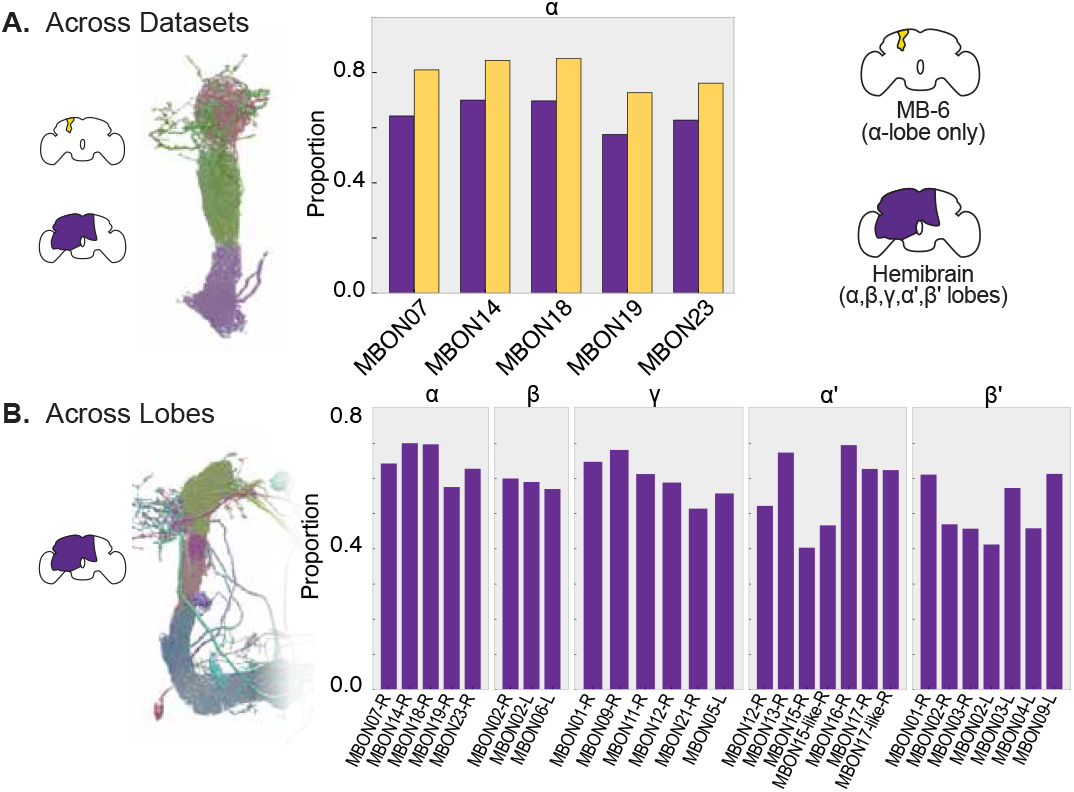
Comparison of convergent synaptic motifs by MBON type. **A**. Proportion of KC-MBON synapses in convergent motifs across datasets; compartment-specific MBONs of the *α* lobe shown **B**. Proportion of KC-MBON synapses in convergent motifs across lobes; compartment-specific MBONs of the *α* and *β* lobe shown. Analysis is limited to MBON cells with at least 100 total synapses.

### 2.3 KC-MBON convergence is prevalent across all MB lobes

Here we asked if convergent motifs occur across all MB lobes. We report the proportion of synapses that participate in convergent motifs for each typical MBONs with dendritic arbors (ipsilateral and/or contralateral) in the MB of the right brain hemisphere of the hemibrain (**Fig. 3**). While the proportion of convergence varies across MBON types, we observed over 50% convergence for each MBON type in the hemibrain.

### 2.4 Newly discovered, multiply convergent motifs are more frequent in some MB lobes than others

During validation, we discovered that some convergent synaptic motifs were tightly spatially clustered. Manual inspection revealed doubly and even triply convergent motifs (**Fig. 4**), where two KC presynapses converge onto more than one MBON neuron. This phenomenon often requires three spatial dimensions to embed while remaining within the 300 nm convergent limit. The double and triple convergence motifs tended to occur in specific cell types and lobes. In very rare cases, we discovered quadruply convergent motifs—an extreme case where two presynaptic sites form connections with four distinct downstream targets. This appears to represent a physical upper limit for convergent motifs. Based on our analysis, we propose that this limitation is not determined by biological or architectural preferences of the neural system, but rather by fundamental spatial constraints within the anatomical microstructure. The physical proximity required for convergent motifs (≤ 300nm) likely restricts the number of distinct postsynaptic targets that can physically access two adjacent presynaptic sites while maintaining proper synaptic architecture.

**Figure 4.**
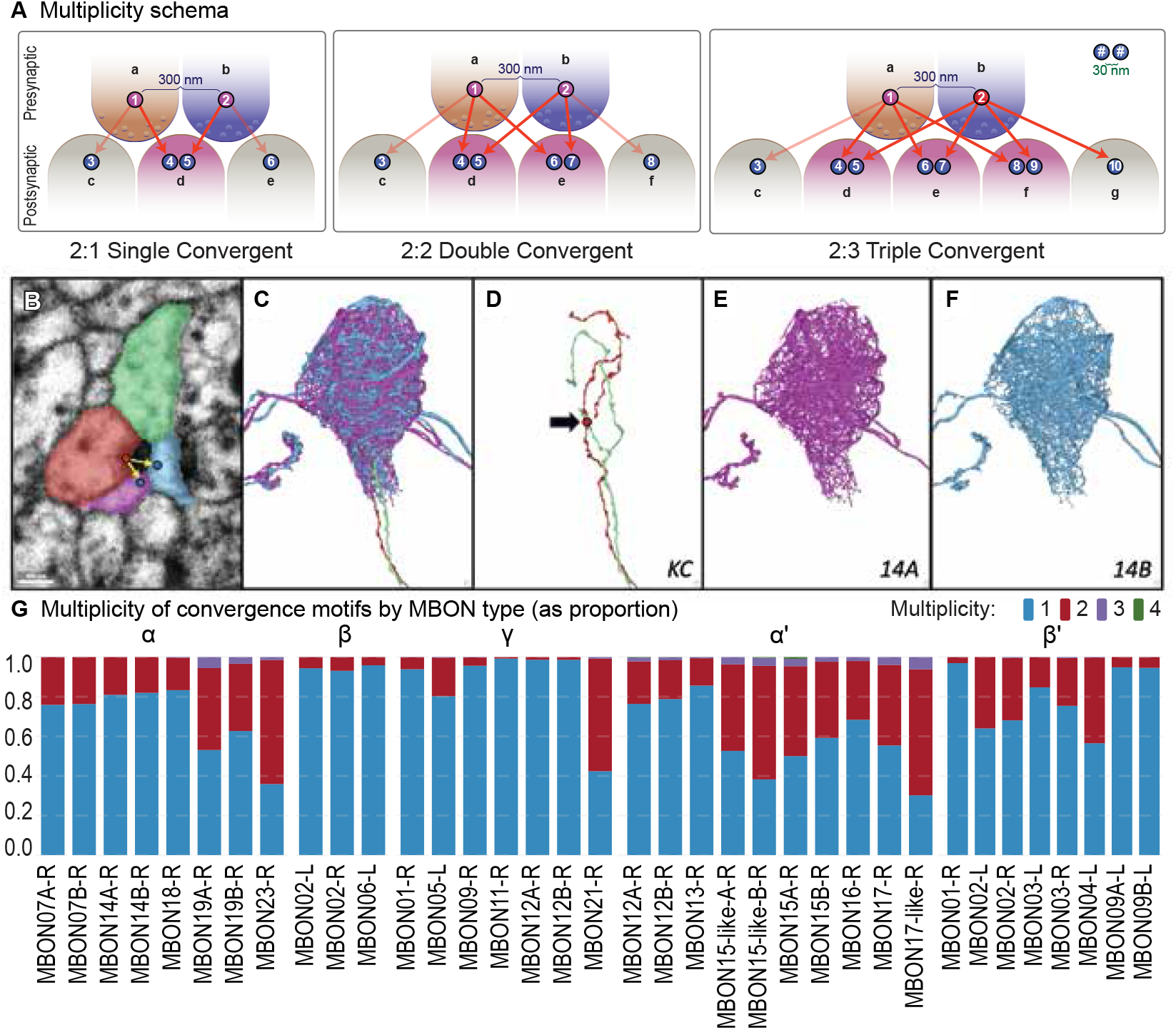
Multiplicity across MBONs. **A**. Schema for single, double, and triple convergent motifs. **B**. Example of KCs and MBONs that form a double motif (panel A, center), seen in electron microscopy. **C**. A 3D reconstruction of the KCs and MBONs from panel B. **D**. The KC participants of the 3D reconstruction from panel C. **E**. The MBON-14A participant of the 3D reconstruction from panel C. **F**. The MBON-14B participant of the 3D reconstruction from panel C. **G**. Proportion of convergent motifs per MBON occurring as single (blue) and multiples (red, purple, green).

We compared the proportion of single versus multiply convergent motifs formed by typical MBONs across different MB lobes in the hemibrain. The MB contains 23 typical and 14 atypical MBON types. Atypical MBONs were recently identified in the hemibrain connectome as MBONs with dendrites within and outside the MB.^17^ The majority of atypical MBONs innervate the *γ* and *β* ‘ lobes. Our analysis focused on multiply convergent motifs involving two or more typical MBONs. Although multiply convergent motifs were observed in all MB lobes, with majority occurring as double motifs, their prevalence was notably lower in the *γ* and *β* lobes (**Fig. 4**). Interestingly, the *γ* lobe—but not the *β* lobe—is also innervated by atypical MBONs. This led us to investigate whether typical MBONs in the *γ* lobe form multiply convergent motifs with atypical MBONs. Our results showed that some typical MBONs, particularly MBON21, frequently participate in multiple motifs. In fact, more than half of the convergent motifs involving MBON21 are multiples (**Fig. S2 supplemental**).

### 2.5 Motif multiplicity is conserved across individuals

Here we ask if the prevalence of multiply convergent motifs is conserved across individuals by comparing the distribution of these motifs per MBON type within the *α*-lobe of MB-6 and hemibrain. Strikingly, with the exception of MBON07, motif multiplicity is highly conserved within MBON types and across brains (**Fig. 5**). For example, within MB-6, 42.7% and 40.6% of convergent motifs formed by MBON19A and MBON19B, respectively, are double motifs, and, within the hemibrain, 19.1% and 18.1% of convergent motifs formed by MBON14A and MBON14B, respectively, are double motifs. More strikingly, more than half, 64.8% in MB-6, and 62.6% in the hemibrain, of convergent motifs formed by MBON23 are double motifs.

**Figure 5.**
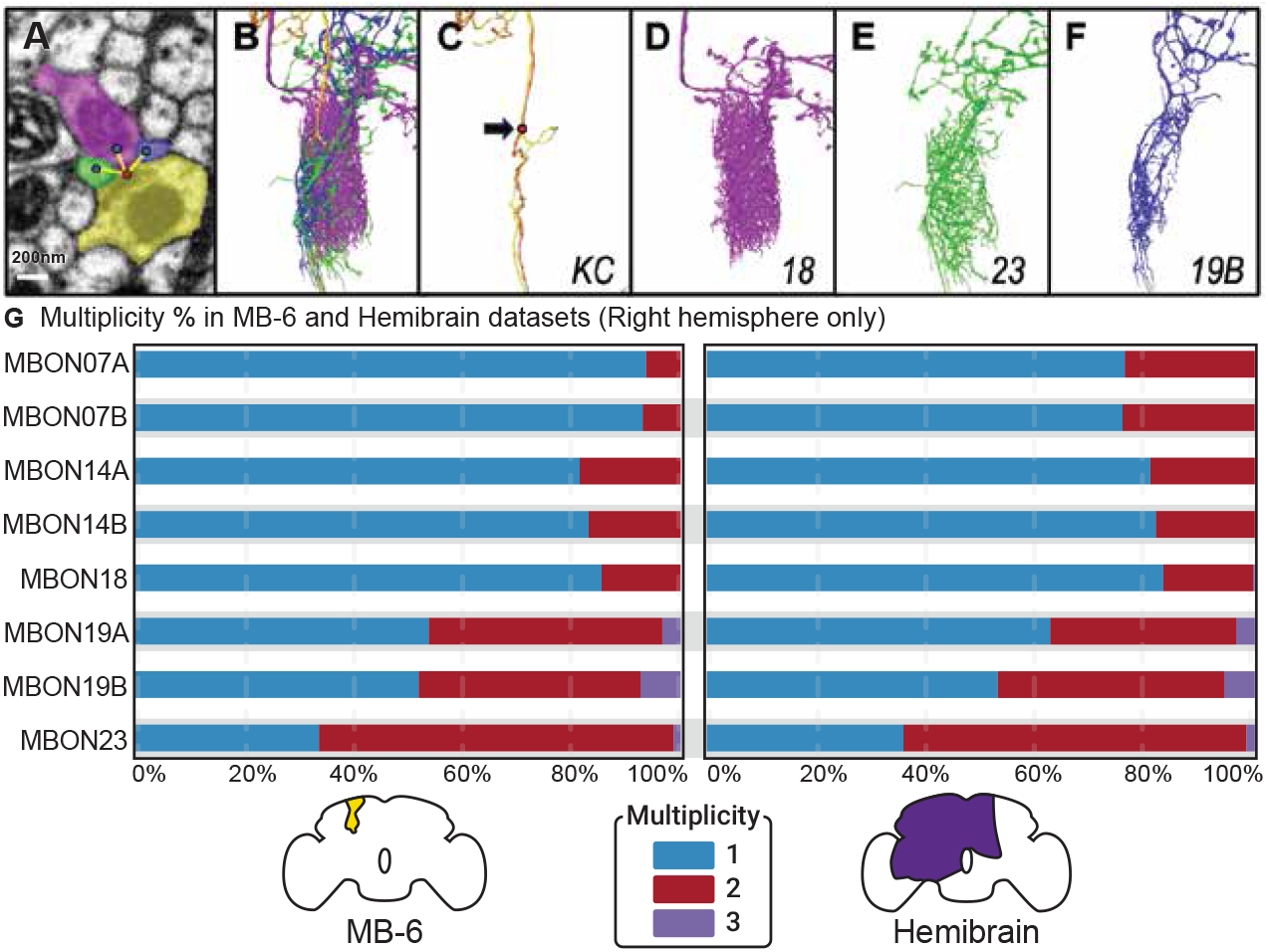
Comparing motif multiplicity across datasets. **A**. Example of KCs and MBONs that form a triple motif in the *α* lobe of the Hemibrain, as seen in electron microscopy, **B**. 3D reconstruction of the KCs and MBONs from panel A, **C**. The KC participants of the 3D reconstruction from panel B, **D-F**. The MBON participants of the 3D reconstruction from panel B, **G**. Proportion of convergent motifs per MBON occurring as single (blue) and multiples (red, purple). See **Table 1** for tabular data and additional commentary.

The striking similarity in motif multiplicity in the *α*-lobe of MB-6 and the hemibrain underscores the conservation and overall consistency of these connectomic datasets, despite minor differences in MBON07 synapse counts (see *Datasets & selection* for additional commentary).

### 2.6 Stereotypic and variable pairing of MBON types at doubly convergent motifs

MBON dendrites divide MB lobes into compartments. Some compartments are innervated by up to four different MBONs. The appearance of doubly convergent motifs prompted us to ask if all MBONs within a compartment can co-occur as post-synaptic targets at double motifs. More specifically, we asked if neurons of the same MBON type can co-occur at double motifs, and if double motifs contain an invariable pair of MBON neurons. To address these questions, for each MBON, we computed the proportion of double motifs formed with other MBONs in the same lobe (**Fig. 6**). We limited our analysis to MBONs that form ≥100 double motifs within a given lobe.

**Figure 6.**
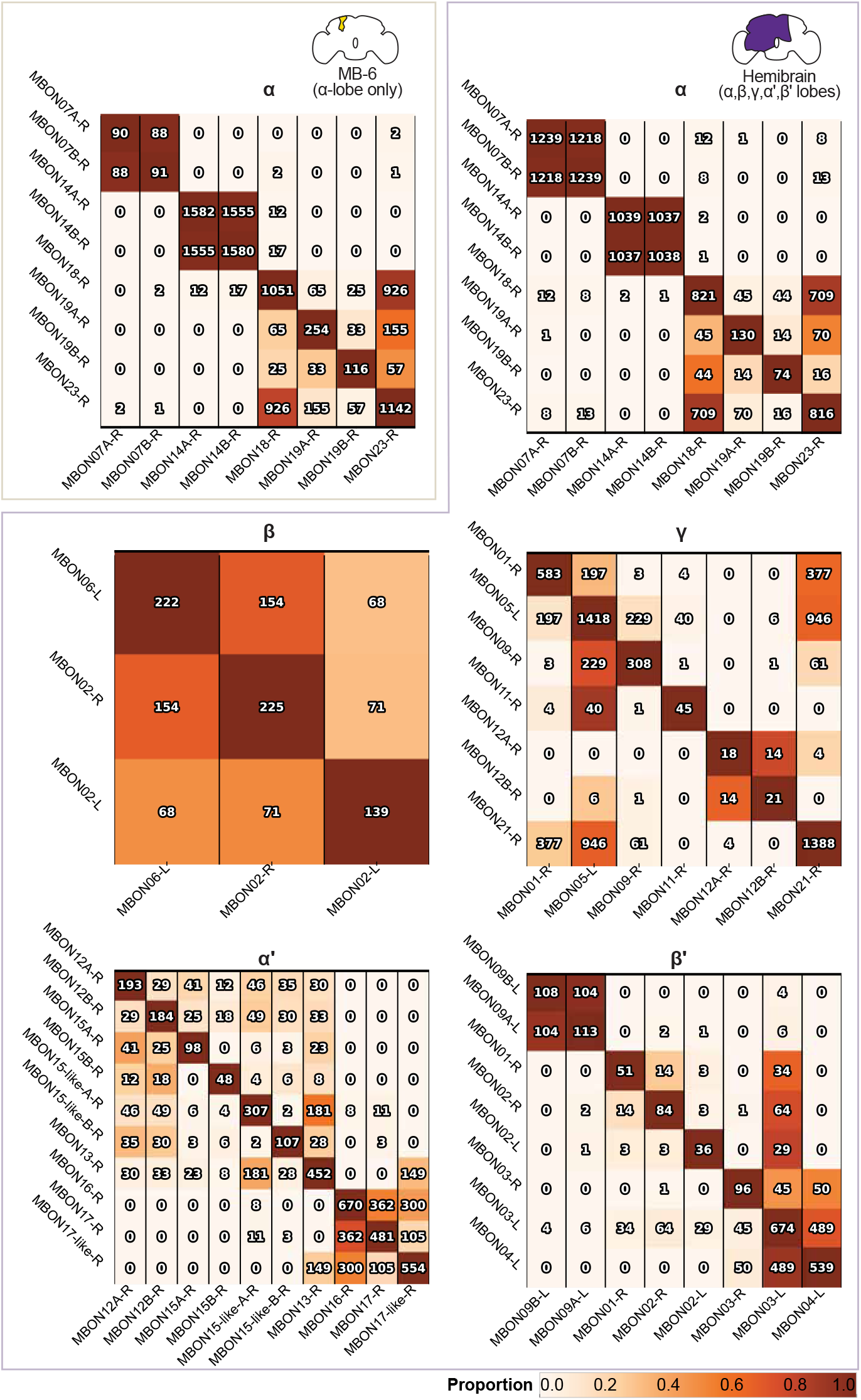
MBON co-occurence. Heatmap showing co-occurrence of MBON neurons at double motifs across lobes in MB-6 and hemibrain. Number of double motifs formed being two different MBONs shown in each square of grid, and total number of double motifs formed by given MBON shown along diagonal.

To address the first question, we analyzed co-occurrence in MBON-07, -09, -12, and -14. Interestingly, in contrast to MBON07A/B, MBON09A/B and MBON14A/B, MBON12A/B rarely co-occur at double motifs (**Fig. 6** *α*’). We also asked if the contralateral and ipsilateral arbors of bilateral neurons of the same MBON type can co-occur at double motifs. For example, MBON02 is a bilateral neuron with an ipsilateral and contralateral dendritic arbor in the *β* 2- and *β* 2’-compartment, respectively. We found that MBON02 neurons in the right and left hemispheres (MBON02-R and -L) co-occur at double motifs in the *β* lobe (**Fig. 6** *β*). Overall, these results highlight the specificity of co-occurrence at double motifs as well as the consistency of reconstruction between ipsilateral and contralateral dendritic arbors within the hemibrain dataset.

Our analysis revealed an additional pattern: double motifs with co-occurring MBONs whose dendrites innervate different but closely apposed compartments. A clear example of this is MBON05, which innervates the *γ*4 compartment (**Fig. 6** *γ*). This MBON forms double motifs with: (1) MBON09 and MBON01, which innervate the flanking compartments *γ*3 and *γ*5 respectively, and (2) MBON21, which spans both *γ*4 and *γ*5 compartments. These findings suggest that double motifs frequently form along compartment boundaries, potentially serving as anatomical markers that more precisely define these functional boundaries. This spatial organization may have implications for information processing between adjacent compartments in the mushroom body.

To determine if double motifs are comprised of an invariable pair of MBON types, we analyzed the double motifs formed by MBON 19A and 19B in both the hemibrain and MB-6 (**Fig. 6** *α*). MBON19A/B form 130 and 74 double motifs, respectively, in the hemibrain, and slightly more, 254 and 116 double motifs, respectively, in MB-6. In both MB-6 and hemibrain, MBON19A and 19B co-occur with each other, as well as MBON18 and 23, in doubly convergent motifs. In MB-6, MBON19A and MBON19B form 49% and 61% of their double motifs with MBON23. However, in the hemibrain, MBON19B forms 54% of its double motifs with MBON23, but MBON19A forms 59% of its double motifs with MBON18. This difference prompted us to look more closely at the synaptic completeness of MBON19A/B in both datasets, where we observed differences in the number of KC-MBON19B synapses per KC: 4.4 in MB-6 and 2.7 in the hemibrain (**Table 1**). This suggests that the postsynaptic completeness (% postsynapses traced to target) is lower for MBON19B in the hemibrain versus MB-6, which may account for the observed differences in double motif partner preferences. Interestingly, this may also explain the differences in visual input to MBON19A and MBON19B observed in the hemibrain.^17^ Excluding the hemibrain’s MBON19B, our findings demonstrate that, while MBON19 can pair with MBON23, MBON18, and each other at double motifs, both MBON19A and MBON19B prefer to partner with MBON23.

### 2.7 Multiply convergent motifs are associated with larger synapses

The discovery of double and triple convergent motifs raised an important question: Do these multiply convergent motifs correspond to physically larger synapses? To investigate this relationship, we used polyadicity—the number of postsynaptic partners per presynapse—as a proxy measure for synapse size. This approach is supported by previous research showing that polyadicity positively correlates with presynaptic membrane area and overall synapse size.^23^ We conducted this analysis on convergent motifs in the *α* lobe, comparing data from both MB-6 and hemibrain datasets.

Our results reveal two key findings: 1. The average polyadicity across all KC-MBON synapses is significantly lower in MB-6 compared to hemibrain (2.55 vs 3.45; Mann-Whitney U test p*<*0.0001).(**Fig. 7A-B**). 2. More importantly, the average polyadicity of double motifs is significantly higher than that of single motifs in both datasets (4.01 and 3.34 in hemibrain; 3.20 and 2.55 in MB-6; Mann-Whitney U test, p*<*0.0001). (**Fig. 7C**).

**Figure 7.**
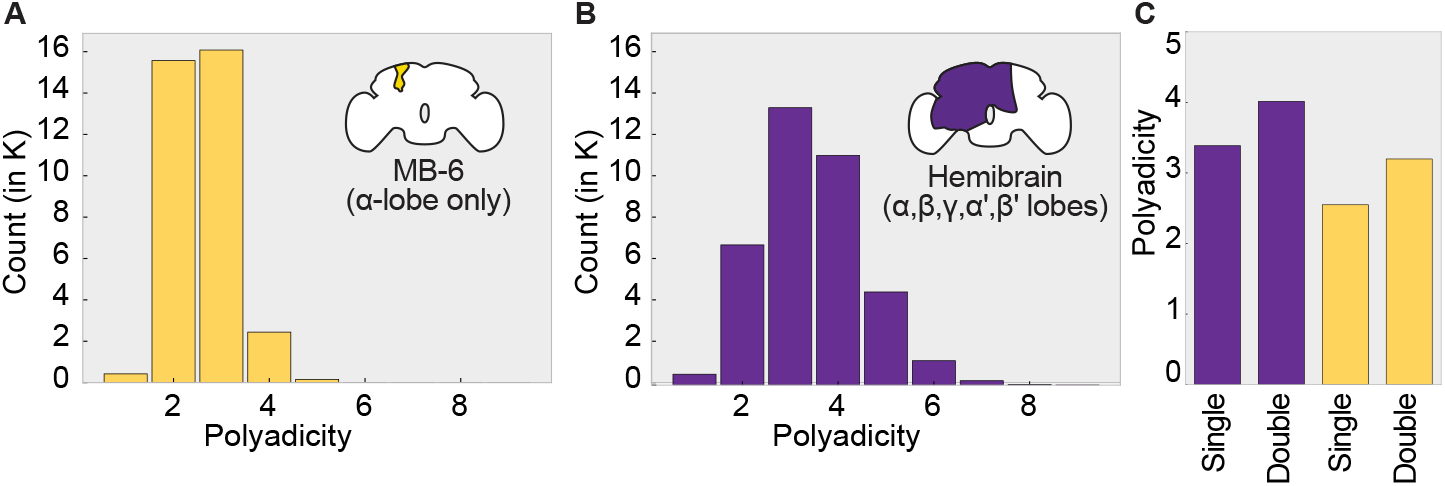
Polyadicity of KC-MBON synapses. **A**. Histogram of KC-MBON polyadicity in MB-6; **B**. Histogram of KC-MBON polyadicity in hemibrain; **C**. Polyadicity of KC-MBON synapses in single versus double motifs in MB-6 and hemibrain Number of double motifs formed being two different MBONs shown in each square of grid, and total number of double motifs formed by given MBON shown along diagonal.

**Figure 8.**
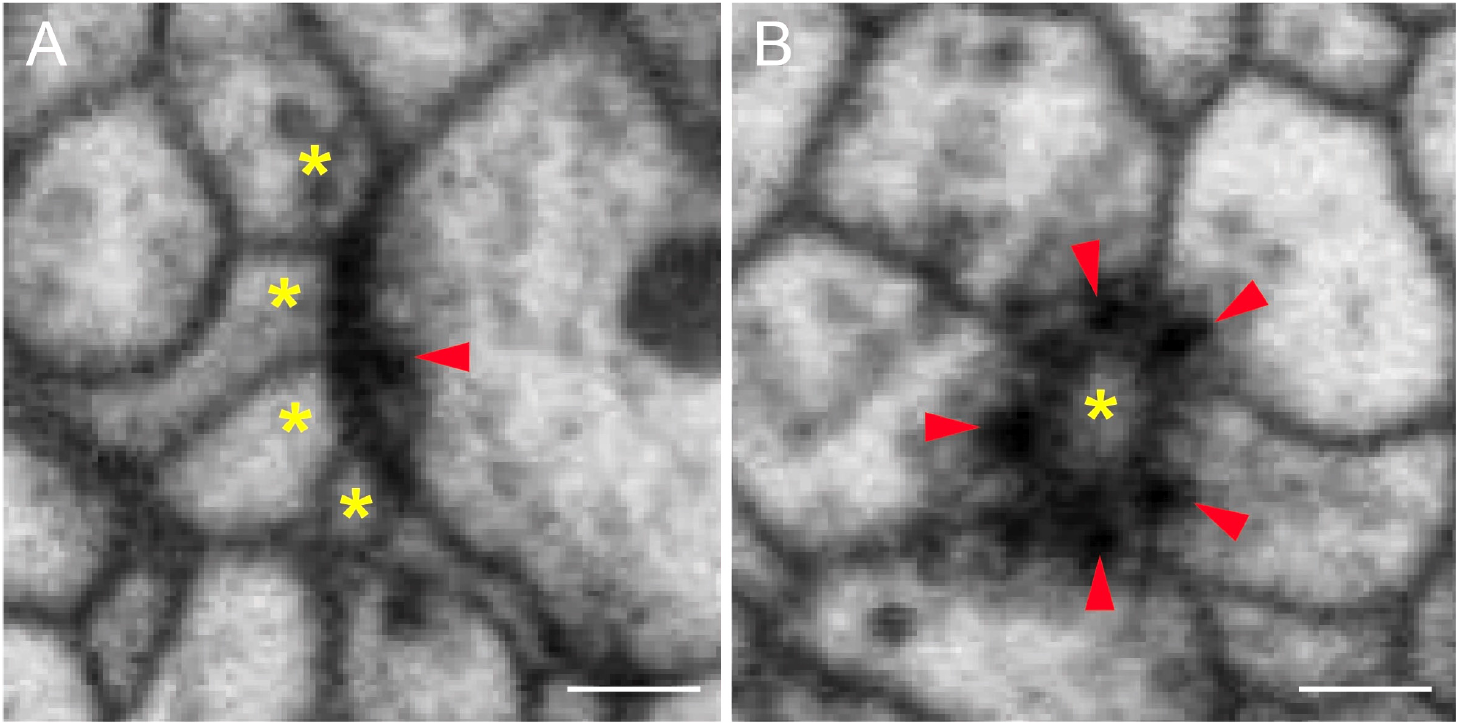
Synaptic configurations in MB-6 as viewed in EM. A. Polyadic, B. Rosette synapse formed by multiple convergent motifs. Presynaptic site (red arrow) and postsynaptic site (*). Scale bar, 100nm.

This second finding strongly supports our hypothesis that multiply convergent motifs (double and triple) are indeed associated with larger synaptic structures. This relationship between convergence multiplicity and synapse size suggests these motifs may represent an efficient structural adaptation for multi-target synaptic communication.

## 3 Discussion

### 3.1 Emerging tools and datasets enable large-scale synaptic motif comparison

To date, the analysis of synaptic motifs across cell types has largely been limited to a few studies, due to the substantial effort required to produce and annotate high-quality EM volumes. Notable examples include polyadic synapses in the Drosophila lamina,^8,9^ clustered synapses in the mouse hippocampus,^24^ polyadic synapses in the Drosophila antennal lobe,^25^ and polyadic synapses in the mouse and rabbit retina.^4^ In particular, less is known about inter-animal variability of synaptic motifs. With the exception of MB-6,^16^ large-scale EM connectomic datasets, though well suited for such analyses, have rarely been leveraged. Using DotMotif, a software tool developed for querying connectome graphs, we were able to conduct a comparative analysis of synaptic motifs across individuals and brain regions. Leveraging two connectomic datasets, MB-6 and the hemibrain, we compared the occurrence of synaptic convergent motifs across cell types (e.g. MBONs), individuals, and subregions of the MB.

The accuracy of any connectome is affected by the quality of synapse detection and synaptic completeness rates (fraction of pre- and post-synapses traced back to reconstructed neurons^1,21^). Both MB-6 and hemibrain connectomes have high synaptic completeness (84–86%) and high synapse detection recall (90%) in the MB lobes, enabling reliable synaptic motif analysis. This study therefore represents the first large-scale comparison of synaptic motifs across individuals and brain subregions (e.g., MB lobes), and demonstrates a proof-of-concept for scalable programmatic motif analysis in emerging, larger connectomes.

### 3.2 Inter-animal variability of synaptic convergent motifs

In agreement with Takemura et al. (2017), we found that the majority of KC-MBON synapses in the *α*-lobe of MB-6 participate in convergent motifs. However, using DotMotif, we observed a slightly lower rate of convergence, 70–85% compared to the 80–93% reported by Takemura et al., depending on the MBON type. This discrepancy may be attributable to methodological differences. Specifically, our initial analysis included false positives when detecting convergent motifs by MBON type. To address this, we revised our approach to detect convergence by specific neuron ID. This refinement likely accounts for the slightly lower percentage of convergence detected by DotMotif relative to previous reports.

Similar to MB-6, we observed over 50% convergence for each MBON type in the hemibrain, demonstrating that synaptic convergence is a general feature of all MB lobes. Interestingly, % convergence per MBON type was slightly lower in the *α* lobe of the hemibrain versus MB-6. Given that both datasets have high synaptic completeness and recall, and that the flies were prepared with similar protocols, we suggest that the observed differences in convergence are largely biological in origin (e.g., sexual dimorphism—MB-6 is male, while the hemibrain is female).

### 3.3 Newly discovered multiply convergent motifs

We hypothesize that the formation of double and triple convergent motifs saves cellular resources and space and is an efficient way to establish new connections or reinforce existing connections, as has been proposed for multi-synaptic spine and bouton formation in the mammalian brain.^14^ An intriguing possibility is that multiple motifs emerge from experience-dependent strengthening of single motifs and their location is a structural signature for learning in the MB lobes.

Using polyadicity as a proxy for synapse size, we demonstrated that synapses involved in double and triple motifs tend to be larger. This finding suggests a potential mechanism for conserving cellular resources—favoring the formation of larger synapses over the addition of multiple smaller ones. Notably, in certain systems such as photoreceptor terminals and the neuromuscular junction, the spatial distribution of presynaptic sites along axons is constrained by nearest-neighbor limitations.^26,27^ In this context, the enlargement of convergent synapses may serve as a structural adaptation to accommodate additional postsynaptic MBON partners, particularly when the formation of new KC presynaptic sites is restricted by these spatial constraints.

### 3.4 Synaptic Exclusion

Synaptic exclusion occurs when repulsive interactions occur between neurites to ensure the correct composition of postsynaptic partners at polyadic synapses. This developmental mechanism has been well characterized in the fly visual system. In the lamina, the first optic lobe neuropil, photoreceptor synapses are consistently composed of a tetrad of postsynaptic elements that invariably include paired L1 and L2 elements.^28,29^ The invariable pairing of L1 and L2 is highly conserved across dipteran species, suggesting this synaptic feature plays a critical role in visual computation in flies.^30^ The occurrence of doubly convergent motifs prompted us to ask whether invariable pairing of MBON types and synaptic exclusion occur in the MB.

Interestingly, we observed that doubly convergent motifs can involve two neurons of the same cell type (e.g., MBON07A and MBON07B), in contrast to photoreceptor synapses in the lamina, where synaptic exclusion mechanisms prevent L1/L1 and L2/L2 pairings. In the lamina, the cell adhesion molecule DSCAM is expressed by L1 neurons and mediates repulsive interactions that underlie this exclusion.^29^ To our knowledge, the expression of DSCAM in MBONs has not been characterized. Our findings suggest that the absence of synaptic exclusion in double motifs may be due to the lack of DSCAM-mediated repulsion.

### 3.5 Limitations of this study

Three datasets containing a partial to complete MB are publicly available: MB-6, Hemibrain, and FlyWire. Our study is limited to MB-6 and Hemibrain. We did not analyze the FlyWire dataset because it did not meet our requirements (see methods) for accurate convergent synapse analysis. In particular, the current FlyWire synapse detection algorithm can not detect many-to-one synaptic configurations, such as convergent motifs, because it assumes a single presynaptic object per detected postsynaptic terminal.^3,31^ Additionally, due to differences in fly rearing, the FlyWire dataset includes extra *γ*KC ^21^ as compared to the Hemibrain. It will be valuable to extend comparisons to whole CNS datasets, when their reconstruction is completed and they are publicly released.

Our observation that MBON19A and MBON19B show inconsistent synapse counts across datasets, unlike other MBON pairs, raises important questions about neural identity and reconstruction quality. To definitively distinguish between incomplete reconstruction versus genuine biological subtypes would require additional analyses beyond the scope of this study. Morphological comparisons of dendritic arbor structure, comprehensive mapping of input connectivity patterns (including non-KC sources), and molecular profiling to identify distinguishing genetic markers would provide crucial evidence. Furthermore, functional characterization through calcium imaging or optogenetic manipulation could reveal whether these neurons exhibit distinct response properties. The existing literature already hints at functional differences, as Li et al. (2020) documented distinct visual input patterns to MBON19A and MBON19B in the hemibrain, supporting the hypothesis that these may represent genuine biological subtypes rather than reconstruction artifacts.

## 4 Methods

### 4.1 Datasets & Selection

The MB of each brain hemisphere contains approximately 2,000 intrinsic neurons called Kenyon Cells (KCs). KC axons run parallel to one another and divide the MB into five lobes: *α, β, α*’, *β* ‘, and *γ*. KCs can be classified into subtypes based on the location of their axons within the lobes. KCab-c,-s,-m, and -p bifurcate and innervate the *α* and *β* lobes. Similarly, KCa’b’-c,-s,-m, and -p innervate the *α*’ and *β* ‘ lobes. KCy-d and -m innervate the *γ* lobe.

MB lobes can be further subdivided into 15 compartmental units by the dendrites of MB output neurons (MBONs; **Table 1**) and the axon terminals of dopaminergic neurons (DANs). To date, 21 DAN types, 20 typical MBON types, and 14 atypical MBON types have been identified.^17^ Additionally, two large neurons, the anterior paired lateral (APL) neuron and the dorsal paired medial (DPM) neuron, arborize throughout the MB lobes, playing a role in global regulation of MB function.

Three connectomic datasets have been generated with electron microscopy to study MB neural circuitry: MB-6, a subvolume containing the *α*-lobe;^16^ Hemibrain, a partial brain volume containing all MB subregions in the right hemisphere;^1,17^ and FlyWire, a whole brain volume containing the MB of both hemispheres.^18^ To serve as a dataset in this study, we require highquality, unbiased synaptic detection that can detect synaptic partners in several configurations of polyadicity and convergence (see 4.2).

A closer examination of MBON07A/B in MB-6 reveals an error in neuPrint, which under-reports the number of MBON07 synapses compared to Takemura et al. (2017). This discrepancy may explain the lower proportion of double motifs detected in MB-6 relative to the hemibrain for this MBON type. Notably, synapse counts for other MBONs in MB-6 are consistent between neuPrint and Takemura et al. (2017). Despite this error in MBON07 synapse counts, the overall consistency of these connectomic datasets makes them well-suited for comparative motif analysis.

To ensure statistical reliability in our convergent motif analyses, we applied a threshold requiring MBONs to have at least 100 total KC-MBON synapses. This threshold was chosen to provide sufficient sample sizes for meaningful statistical comparisons of motif patterns across different MBON types and between datasets. The few MBONs with fewer than 100 synapses were excluded from quantitative analyses to avoid spurious results that could arise from small sample sizes.

### 4.2 Synapse Detection and Motifs in Drosophila

Synapses can be visually detected in Drosophila by the presence of an electron-dense projection called the T-bar.^8^ To date, the majority of synapses studied in the fly brain are polyadic (**Fig. 8A**), where a single T-bar has multiple postsynaptic partners.^20^ Fly synapse detection has therefore been optimized to detect one-to-one and one-to-many synaptic connections. In MB-6 and hemibrain, synapse detection was additionally optimized to detect convergent motifs (defined in 4.3; **Fig. 8B**), particularly in the MB, where many-to-one synaptic arrangements have been commonly observed.^1,16^ Unlike MB-6 and hemibrain, FlyWire’s synapse detection method was less suited to detect many-to-one synapses,^3^ and thus could not be used in this study.

### 4.3 Defining convergence

We first looked to the existing literature for a formal definition of synaptic convergence. Scheffer et al. (^1^) define convergence as a tight grouping of two axons whose active zones are less than 300 nm apart and are presynaptic to the same target. These many-to-one motifs (**Fig. 1**) are well-established in the invertebrate literature (locust,^32^ insect,^33^ octopus,^34^ fruit fly^16^).

We validated this definition by analyzing the distribution of distances between presynaptic sites that share a common postsynaptic target (see Fig. S1 in Supplemental). Our analysis confirmed that 300 nm represents an appropriate threshold, as it captures the majority of biologically relevant convergent connections while excluding more distant, potentially unrelated synaptic pairs.

### 4.4 Large-scale detection of convergent motifs with DotMotif

Here we employed DotMotif, a standard network motif search tool^2^ for analysis of convergent synaptic motifs within the hemibrain and MB-6 datasets. To our knowledge, this represents the largest-scale analysis of a synaptic motif across any connectome to date. We allowed for KCs of different types (KCab-c and KCab-m) to converge onto the same MBON target.

We evaluated the accuracy of our convergent motif detection in the hemibrain by manually inspecting approximately 100 motif predictions across 3 lobes, *α, β* and *γ* using the Neuroglancer viewer in neuPrint, which has been optimized to visualize individual presynaptic sites and their associated postsynaptic connections. Although nearly all motif predictions were correct, we uncovered a few interesting false and true positives that prompted us to refine our motif detection. More specifically, we uncovered false positive motifs where each KC was presynaptic to the same MBON type but different neuron IDs. In addition, we uncovered false positive motifs where both KCs were presynaptic to the same MBON neuron ID, but the presynapses were located on the same KC neuron ID. The latter arrangement resembles the clustered, multiple synapse arrangement observed in mammalian brains, where one axon forms multiple, closely spaced synapses, with the same dendritic branch (ref).

During our manual inspection, we identified an intriguing pattern: some true positive motifs showed both KCs converging onto multiple MBON neuron IDs simultaneously. This observation led us to conduct a thorough examination of these “multiply convergent” motifs. We manually inspected all 85 additional instances of these multiple-motifs in the *α*2-compartment, a region where the dendritic arbors of MBON 18, 19, and 23 overlap. We hypothesized that multiply convergent motifs would be most prevalent in compartments where multiple MBONs have overlapping dendritic arbors. Our inspection confirmed this hypothesis, revealing double and triple convergent motifs involving various combinations of MBON 18, 19A, 19B, and 23 in the *α*2 compartment. These findings prompted us to conduct a comprehensive analysis of the distribution of both single and multiply convergent motifs across all MBONs, lobes, and compartments (see section on convergent motif multiplicity).

## 5 Acknowledgements

The authors thank Daniel Xenes and Hannah Martinez for their technical guidance, and Stuart Berg for assistance with the MB-6 dataset.

We gratefully acknowledge the support of Johns Hopkins Applied Physics Laboratory internal research and development funding. Research reported in this publication was supported by the National Institute Of Mental Health of the National Institutes of Health under Award Number R24MH114785. The content is solely the responsibility of the authors and does not necessarily represent the official views of the National Institutes of Health.

## 6 Data Availability

The datasets analyzed in this work are publicly accessible from neuPrint (https://neuprint.janelia.org/?dataset=hemibrain%3Av1.2.1) and BossDB (https://bossdb.org).

All remaining data and analyses produced in this work will be made available upon publication and upon request.

## 7. Supplemental

### 7.1 Convergence distance thresholds

We illustrate in **Fig. S1** the distribution of synaptic distances for pairs of neurons with the same downstream target neuron. This analysis validates our use of the 300nm threshold for defining convergent motifs, as the frequency distribution shows a clear peak at shorter distances and decreases rapidly beyond 300nm. The red line marking this threshold demonstrates that it effectively captures the majority of biologically relevant convergent connections while excluding more distant, potentially unrelated synaptic pairs.

**Figure S1.**
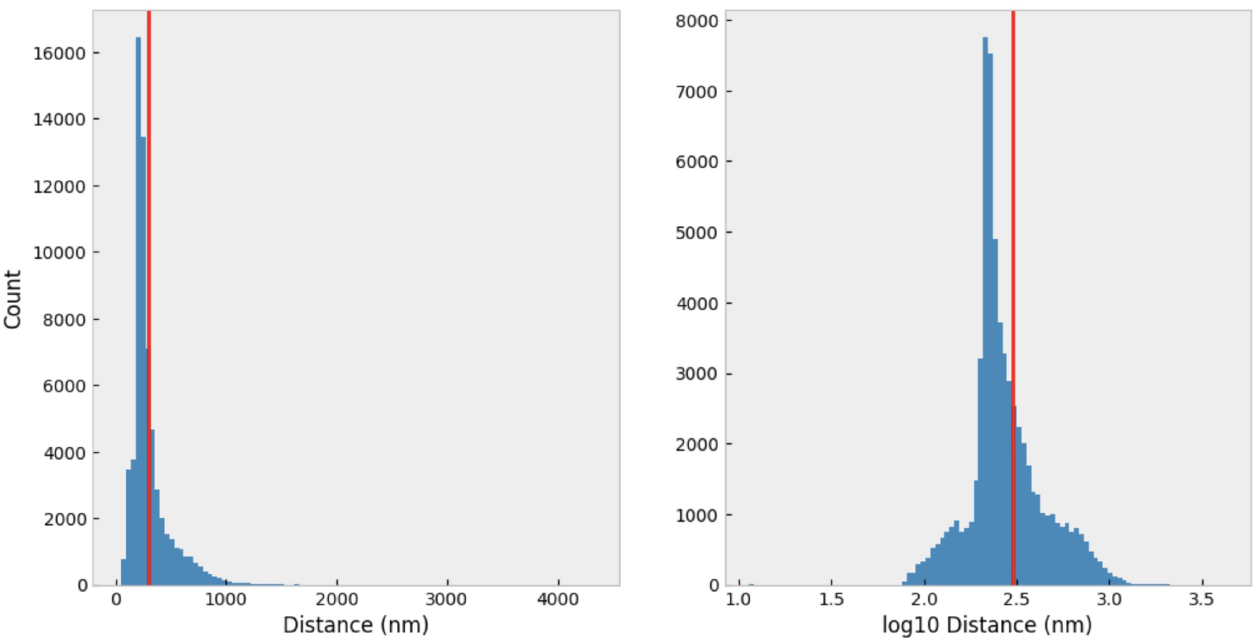
A histogram of synaptic distances for upstream targets that synapse on the same downstream neuron. The accepted convergence threshold of 300nm is drawn in red.

### 7.2 Typical MBONs form mulitply convergent motifs with atypical MBONs in *γ* lobe of hemibrain

The *γ* lobe is innervated by seven typical MBONs (01, 05, 09, 11, 12A, 12B, 21) and ten atypical MBONs (20, 24, 25, 27, 29, 30, 32, 33, 34, 35). Here we computed the proportion of single, double, triple, and quadruple motifs formed by each typical MBONs with atypical MBONs (**Fig. S2**).

**Figure S2.**
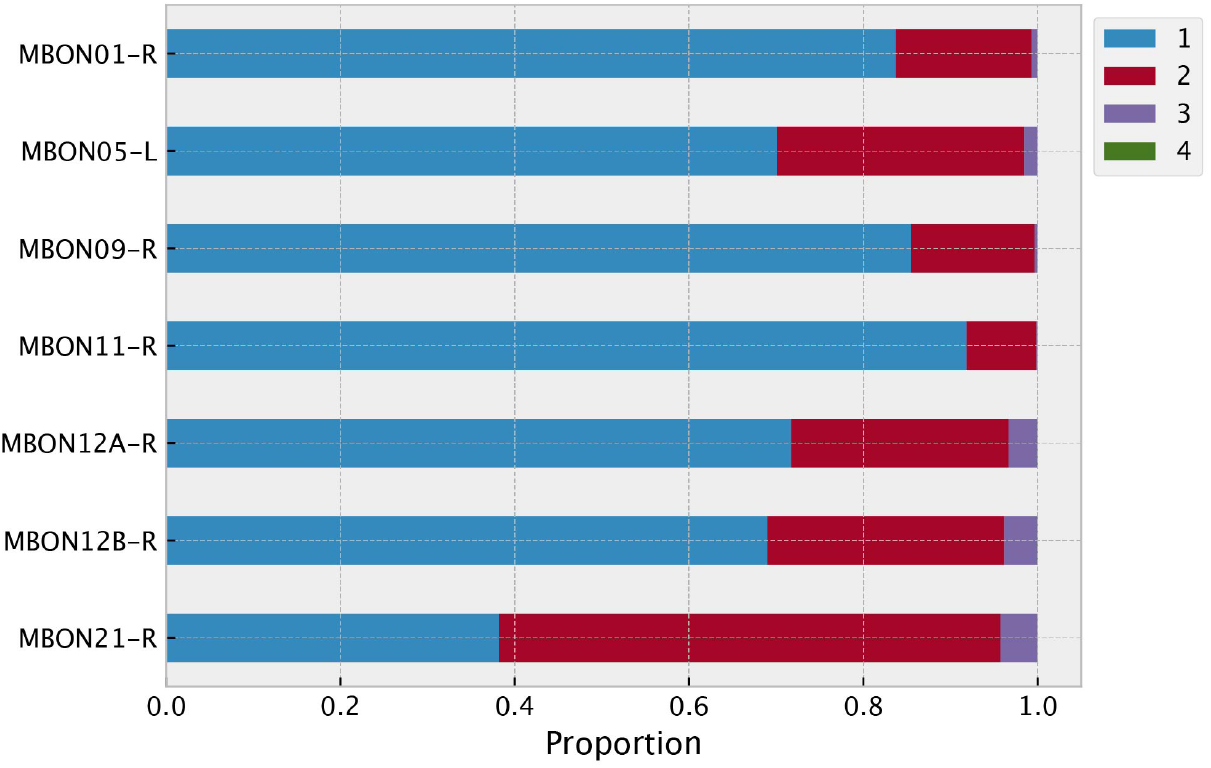
Motif multiplicity across typical MBONs in *γ* lobe. Proportion of convergent motifs per typical MBON occurring as single (blue) and multiples (red, purple, green)

## References

1. Scheffer, L. K. et al. A connectome and analysis of the adult drosophila central brain. 9, e57443. Publisher: eLife Sciences Publications, Ltd.

2. Matelsky, J. K. et al. DotMotif: an open-source tool for connectome subgraph isomorphism search and graph queries. 11, DOI: 10.1038/s41598-021-91025-5. Publisher: Springer Science and Business Media LLC.

3. Buhmann, J. et al. Automatic detection of synaptic partners in a whole-brain drosophila electron microscopy data set. 18, 771–774, DOI: 10.1038/s41592-021-01183-7. Publisher: Nature Publishing Group.

4. Yu, W.-Q. et al. Distinctive synaptic structural motifs link excitatory retinal interneurons to diverse postsynaptic partner types. Cell reports 42 (2023).

5. Gruber, L. et al. The unique synaptic circuitry of specialized olfactory glomeruli in drosophila melanogaster. bioRxiv 2022–09 (2022).

6. Kasthuri, N. et al. Saturated reconstruction of a volume of neocortex. 162, 648–661, DOI: 10.1016/j.cell.2015.06.054.

7. Hwang, Y.-S. Studies on the 3d ultrastructure of synaptic inputs to distinct GABAergic neurons in the primary visual cortex.

8. Meinertzhagen, I. A. & O’neil, S. Synaptic organization of columnar elements in the lamina of the wild type in drosophila melanogaster. Journal comparative neurology 305, 232–263 (1991).

9. Meinertzhagen, I. & Sorra, K. Synaptic organization in the fly’s optic lamina: few cells, many synapses and divergent microcircuits. Progress brain research 131, 53–69 (2001).

10. Matelsky, J. K. et al. Data-driven motif discovery in biological neural networks. bioRxiv 2023–10 (2023).

11. Homberg, U. & Muller, M. Ultrastructure of GABA- and Tachykinin-Immunoreactive Neurons in the Lower Division of the Central Body of the Desert Locust. Frontiers Behavioral Neuroscience 10, 230, DOI: 10.3389/fnbeh.2016.00230 (2016).

12. Hulse, B. K. et al. A connectome of the Drosophila central complex reveals network motifs suitable for flexible navigation and context-dependent action selection. eLife 10, e66039, DOI: 10.7554/elife.66039 (2021).

13. Nikonenko, I., Jourdain, P. & Muller, D. Presynaptic remodeling contributes to activity-dependent synaptogenesis. Journal Neuroscience 23, 8498–8505 (2003).

14. Yang, Y., Lu, J. & Zuo, Y. Changes of synaptic structures associated with learning, memory and diseases. Brain Science Advances 4, 99–117 (2018).

15. Giese, K. P., Aziz, W., Kraev, I. & Stewart, M. G. Generation of multi-innervated dendritic spines as a novel mechanism of long-term memory formation. Neurobiol. learning memory 124, 48–51 (2015).

16. Takemura, S.-y. et al. A connectome of a learning and memory center in the adult drosophila brain. 6, e26975, DOI: 10.7554/eLife.26975. Publisher: eLife Sciences Publications, Ltd.

17. Li, F. et al. The connectome of the adult drosophila mushroom body provides insights into function. 9, e62576, DOI: 10.7554/eLife.62576. Publisher: eLife Sciences Publications, Ltd.

18. Dorkenwald, S. et al. Neuronal wiring diagram of an adult brain. 2023.06.27.546656, DOI: 10.1101/2023.06.27.546656.

19. Aso, Y. et al. The neuronal architecture of the mushroom body provides a logic for associative learning. 3, e04577, DOI: 10.7554/eLife.04577. Publisher: eLife Sciences Publications, Ltd.

20. Prokop, A. & Meinertzhagen, I. A. Development and structure of synaptic contacts in drosophila. In Seminars in cell & developmental biology, vol. 17, 20–30 (Elsevier, 2006).

21. Schlegel, P. et al. Whole-brain annotation and multi-connectome cell typing of drosophila. Nature 634, 139–152 (2024).

22. Lu, Z. et al. En bloc preparation of Drosophila brains enables high-throughput FIB-SEM connectomics. Frontiers Neural Circuits 16, 917251, DOI: 10.3389/fncir.2022.917251 (2022).

23. Butcher, N. J., Friedrich, A. B., Lu, Z., Tanimoto, H. & Meinertzhagen, I. A. Different classes of input and output neurons reveal new features in microglomeruli of the adult Drosophila mushroom body calyx. Journal Comparative Neurol. 520, 2185–2201, DOI: 10.1002/cne.23037 (2012).

24. Bloss, E. B. et al. Single excitatory axons form clustered synapses onto CA1 pyramidal cell dendrites. Nature Neuroscience 21, 353–363, DOI: 10.1038/s41593-018-0084-6 (2018).

25. Gruber, L. et al. The unique synaptic circuitry of specialized olfactory glomeruli in Drosophila melanogaster. eLife DOI: 10.7554/elife.88824 (2023).

26. Meinertzhagen, I. A. & Hu, X. Evidence for site selection during synaptogenesis: The surface distribution of synaptic sites in photoreceptor terminals of the fliesMusca andDrosophila. Cellular Molecular Neurobiol. 16, 677–698, DOI: 10.1007/bf02151904 (1996).

27. Meinertzhagen, I., Govind, C., Stewart, B., Carter, J. & Atwood, H. Regulated spacing of synapses and presynaptic active zones at larval neuromuscular junctions in different genotypes of the flies Drosophila and Sarcophaga. Journal Comparative Neurol. 393, 482–492, DOI: 10.1002/(sici)1096-9861(19980420)393:4⟨482::aid-cne7⟩3.0.co;2-x (1998).

28. Frohlich, A. & Meinertzhagen, I. Quantitative features of synapse formation in the fly’s visual system. I. The presynaptic photoreceptor terminal. The Journal Neuroscience 3, 2336–2349, DOI: 10.1523/jneurosci.03-11-02336.1983 (1983).

29. Millard, S. S., Lu, Z., Zipursky, S. L. & Meinertzhagen, I. A. Drosophila Dscam Proteins Regulate Postsynaptic Specificity at Multiple-Contact Synapses. Neuron 67, 761–768, DOI: 10.1016/j.neuron.2010.08.030 (2010).

30. Shaw, S. R. & Meinertzhagen, I. A. Evolutionary progression at synaptic connections made by identified homologous neurones. Proceedings National Academy Sciences 83, 7961–7965, DOI: 10.1073/pnas.83.20.7961 (1986).

31. Yu, S.-c. et al. New synapse detection in the whole-brain connectome of drosophila. bioRxiv 2025–07 (2025).

32. Rind, F. C. & Simmons, P. J. Local circuit for the computation of object approach by an identified visual neuron in the locust. Journal Comparative Neurol. 395, 405–415 (1998).

33. Schurmann, F.-W. Fine structure of synaptic sites and circuits in mushroom bodies of insect brains. Arthropod Structure & Development 45, 399–421 (2016).

34. Bidel, F. et al. Connectomics of the octopus vulgaris vertical lobe provides insight into conserved and novel principles of a memory acquisition network. Elife 12, e84257 (2023).

